# Field-Correcting GRAPPA (FCG): a technique to correct spatiotemporal-varying phase errors in Echo Planar Imaging

**DOI:** 10.64898/2026.07.10.737642

**Authors:** Nan Wang, Daniel Abraham, Zachary Shah, Yimeng Lin, Xiaozhi Cao, Hua Wu, Jonathan Polimeni, Renzo Huber, Qiang Liu, Lipeng Ning, Yogesh Rathi, Carl-Fredrik Westin, Hendrik Mattern, Oliver Speck, Baolian Yang, Nastaren Abad, Congyu Liao, Adam B. Kerr, Kawin Setsompop

## Abstract

**Purpose:** To develop a Field-Correcting GRAPPA (FCG) technique to correct the spatiotemporal-varying phase errors in EPI caused by eddy currents.

**Methods:** The fast-changing gradient in EPI causes strong eddy current effects and associated spatiotemporal-varying phase errors, producing significant image artifacts. The use of higher gradient amplitude, slew rate, and ramp sampling factor for faster imaging exacerbates this problem. In this work, FCG was developed to address this challenge by using a multi-layer perceptron (MLP) to provide a compact representation of a family of GRAPPA-like kernels that correct the spatiotemporal-varying phase errors in the data. A dedicated calibration pipeline was designed to acquire high-quality source and target data for MLP training in both slice-by-slice and simultaneous multi-slice (SMS) acquisitions. To validate FCG’s assumptions and performance, a field camera was used to provide ground-truth measurement of phase patterns. The performance of FCG was further validated on phantom and in vivo experiments using demanding EPI trajectories across multiple 3T and 7T systems.

**Results:** Field camera measurements revealed strong spatiotemporal phase variations along the *k_x_* direction that repeat along ky during EPI readouts. The experiments on high-performance systems across 3T and 7T demonstrate that FCG can provide superior correction for the artifacts induced by spatiotemporal-varying phase errors compared with existing approaches.

**Conclusion:** FCG is an effective and robust method for correcting spatiotemporal phase errors in EPI, enabling improved image quality on high-performance systems.

## 1 INTRODUCTION

Echo Planar Imaging (EPI)^1^ is a fast and efficient sampling strategy in MRI. It has been widely used to acquire images with diverse contrast (diffusion weighted images ^2,3^, multi-parametric and susceptibility weighted mapping ^4,5^, perfusion ^6^, fMRI^7,8^, etc) with applications across a broad range of anatomical regions from the brain to peripheral organs ^9^.

Nonetheless, EPI is prone to acquisition artifacts, owning to its rapid k-space traversal using fast-changing, polarity-switching gradient encoding. Gradient errors from multiple sources, including delay^10^, non-linearity^11^, and eddy current^12,13^, typically manifest as low-spatial-order phase errors between odd (RO+) and even (RO-) frequency encoding lines (*k*_*x*_), leading to “Nyquist ghost”^14^ that can significantly degrade image quality. A common strategy to improve the quality of the reconstruction is to perform linear (constant plus first-order) phase correction along the frequency encoding direction. The coefficients for both constant and first order terms are nearly flipped sign between RO+ and RO-readouts. These linear phase correction coefficients are typically estimated using a set of back-and-forth 1D reference kx lines at center k-space (ky=kz=0) at the beginning of the scan ^15,16^, and then applied to the actual time-series EPI data. This method, known as Linear Ghost Correction (LGC), has become a standard correction step in EPI and substantially improves image quality in many applications.

However, LGC does not address higher-order (non-linear) phase errors, which can lead to residual ghosting. To mitigate these effects, various approaches have been developed. A category of methods acquires each *k*_*x*_ line in EPI data twice with opposing polarities in either consecutive echoes ^17^ or separate acquisitions ^18,19,20^ and averages them to suppress ghosting artifacts, albeit at the cost of halving the temporal resolution or doubling the scan time. To overcome this issue, Dual-Polarity Grappa (DPG)^21,22^ was developed and has now been widely adopted. Instead of acquiring all imaging data with dual polarities, DPG acquires only the calibration data with dual polarities, synthesizes corresponding ghost-free targets by averaging the dual-polarity calibration data, and trains a pair of GRAPPA kernels that map the corrupted RO+ or RO-data to the ghost-free data, respectively. During imaging, single-average time-series EPI data are corrected using these pre-trained kernels.

Although DPG effectively mitigates nonlinear phase errors in many applications, it has inherent limitations. Specifically, the use of a single GRAPPA kernel for all the RO+ or RO-readouts neglects spatiotemporal variations of phase errors along the readout direction. Such spatiotemporal-varying phase errors becomes increasingly problematic nowadays because: (1) modern high-performance gradient systems enable higher gradient amplitudes and slew rates, leading to stronger and more complex eddy currents; (2) the use of large ramp sampling factors to maximize sampling efficiency under gradient constraints further amplifies these effects; (3) the adoption of high in-plane and slice accelerations, aimed at improving spatial resolution and reducing distortion or acquisition time, increases the sensitivity of image reconstruction to such phase errors. Recent work has shown that, with submillimeter spatial resolution and high undersampling rate, the spatiotemporal-varying phase errors cause severe ghosting and shading artifacts ^20^. In these regimes, DPG may be insufficient, whereas the direct dual-polarity averaging (DPA) of EPI data acquired with dual polarities in two consecutive acquisitions can still produce high-quality images, at a cost of doubled scan time.

To address the spatiotemporal-varying phase errors in high-performance EPI without increasing scan time, we developed the Field-Correcting GRAPPA (FCG) technique. FCG is a kernel-based approach that requires the same calibration data as for DPG. Instead of training only one pair of GRAPPA kernels, FCG employs a multi-layer perceptron (MLP) to compactly represent a family of kernels capable of correcting spatiotemporal field variations along *k*_*x*_. A practical data correction pipeline was developed to facilitate robust application of FCG across a wide range of EPI protocols. The proposed method was evaluated on demanding EPI trajectories with various acceleration schemes, including in-plane acceleration and simultaneous multi-slice (SMS) imaging ^23,24^, across multiple high-performance systems (whole-body and head-only) at both 3T and 7T. Phantom and in vivo experiments demonstrate that FCG provides improved artifact suppression and image quality compared with existing approaches.

## 2 THEORY

### 2.1 Nonlinear phase errors during image encoding

During MRI image encoding, the acquired signal *S*(*t*) at time point *t* can be expressed as

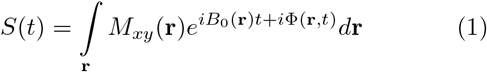

where *M*_*xy*_(**r**) denotes the transverse magnetization at spatial location **r** = (*x, y, z*), *B*_0_(**r**) represents the main field inhomogeneity, and Φ(**r**, *t*) captures both the encoding trajectory driven by the imaging gradients **k**(*t*) and additional phase contributions arising from system imperfections. Eddy currents are a major source of these phase errors.

Field monitoring devices can be used to characterize these phase errors during data acquisition ^25^. For example, Skope (Skope MR Technologies Inc., Zurich, Switzerland), a commercial field camera, enables direct measurement of the spatiotemporal phase evolution and its decomposition into third-order spherical harmonic basis functions with corresponding time-varying coefficients:

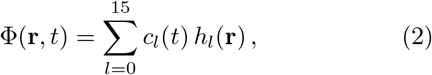

where *h*_*l*_(**r**) denotes the *l*th real-valued spherical harmonic basis function, and *c*_*l*_(*t*) is the corresponding temporal coefficient. The measured phase information can be incorporated into image reconstruction to mitigate phase inconsistencies and produce ghost-free images.

### 2.2 Existing approaches to correct nonlinear phase errors: Dual-polarity averaging (DPA)

Field cameras are expensive and require substantial effort for setup, measurement, and reconstruction, which limits their widespread adoption. In practice, dual-polarity averaging (DPA) provides a robust and practical approach for nearly ghost-free outcomes. The phase term Φ(**r**, *t*) in Equation 1 can be expressed as:

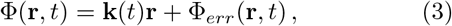

where **k**(*t*) = (*k*_*x*_(*t*), *k*_*y*_(*t*), *k*_*z*_(*t*)) denotes the k-space trajectory, and Φ_*err*_(**r**, *t*) represents phase perturbations, which includes the gradient delay, eddy-current–induced phase errors, the difference of *B*_0_ phase accumulations between RO+ and RO-readouts, etc. By reparameterizing Φ_*err*_(*t*) in terms of **k**, the signal equation (Equation 1), while temporarily neglecting *B*_0_ term, can be written as:

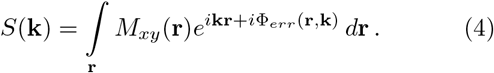

The eddy-current-induced phase is driven by the time derivative of the gradient waveform. With reversed gradient polarity, a dominant portion of Φ_*err*_(**r, k**) is commonly assumed to be antisymmetric. Accordingly, the RO+ and RO-signals can be expressed as:

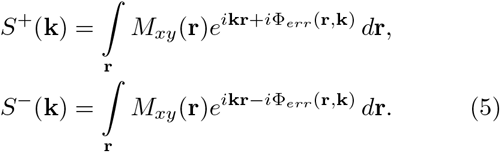

Averaging *S*^+^ and *S*^−^ yields:

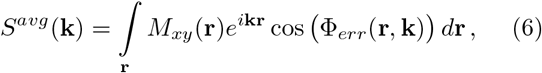

which forms the basis of DPA. For Equation 6 to produce high-quality images, two conditions must be satisfied: (1) Φ_*err*_ is sufficiently small such that cos(Φ_*err*_) ≈ 1, otherwise amplitude modulation is introduced along the EPI trajectory; (2) the k-space trajectory **k** is identical for RO+ and RO-data (Equations 5 and 6), which typically requires linear gradient error correction, aka LGC, before averaging.

In practice, due to the complex interactions between time-varying magnetic fields and hardware, the eddy-current–induced phase is not perfectly antisymmetric between RO+ and RO-. Nevertheless, because the dominant phase components are approximately antisymmetric after removing gradient delay and other linear terms, DPA remains effective in suppressing artifacts. Note that the *B*_0_ phase accumulation along the *k*_*x*_ that are anti-symmetric for RO+ and RO-lines can also be mitigated by DPA with additional details provided in Supporting Information Section A.

### 2.3 Existing approaches to correct nonlinear phase errors: Dual-Polarity GRAPPA (DPG)

Unlike DPA, which doubles the scan time for every repetition, Dual-Polarity GRAPPA (DPG)^21,22^ only acquires a set of fully-sampled calibration data in dual polarities, which serve both phase-error correction and parallel-imaging reconstruction. In conventional DPG, multiple kernels are trained to perform phase-error correction and parallel-imaging reconstruction jointly ^21,22^. In recent years, to allow flexible choice of correction approaches and parallel-imaging protocols, DPG can also refer to only the phase-error correction step that is performed prior to the standard or customized parallel-imaging reconstruction ^26^. In this work, DPG refers to the phase-error correction only.

The calibration data acquisition and training data preparation are shown in Figure 1. The calibration data are acquired with the same EPI trajectory as for imaging acquisition to ensure consistent eddy current behavior and geometric distortion. In the scenario of segmented EPI trajectories with *M* segments, data for each polarity acquire *M* interleaves to form a fully-sampled calibration data, resulting in 2*M* shots for calibration in dual polarities. The set of all the RO+ readouts from dual-polarity calibration data forms *C*^+^, while the set of all RO-readouts forms *C*^−^. Following Equation 6, the complex average of *C*^+^ and *C*^−^ forms ghost-free target data *C*^*avg*^. A pair of GRAPPA kernels, *G*^+^ and *G*^−^, are then trained to map the polarity-specific data to the ghost-free target:

**FIGURE 1.**
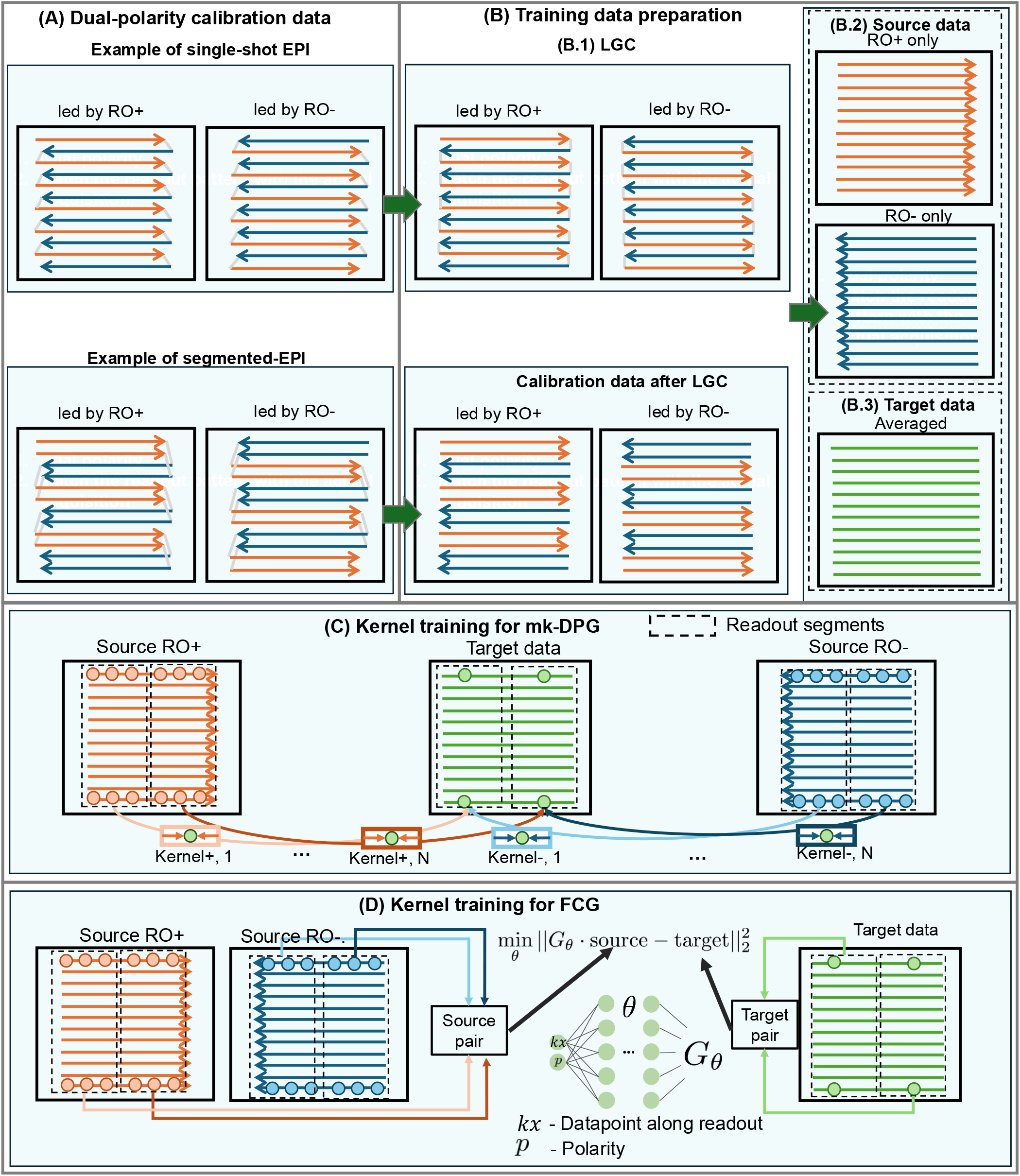
Fig 1: (A) Scheme to acquire dual-polarity calibration data. This is the same as what have been specified in DPG. The calibration data uses the same EPI trajectories as for time-series acquisition. In the scenario with segmented EPI, multiple interleaves are acquired to fulfill a fully-sampled calibration. (B) Training data preparation. The dual-polarity calibration data are corrected with linear-ghost correction (LGC). Then all the RO+ readouts are taken to form a complete set of RO+ only data. The RO-only data are formed analogously. The complex average of RO+ only and RO-only data form the ghost-free target data. Kernels are then trained to map the corrupted to the ghost-free k-space. (C) Kernel training for mk-DPG. The source and target data are divided into N segments along the readout direction. For each segment a pair of positive and negative kernels are trained, resulting in N pairs of kernels. (D) Kernel training for FCG. FCG prepares many source and target data pairs along kx. Instead of training them separately as in mk-DPG, FCG uses a MLP network to joint

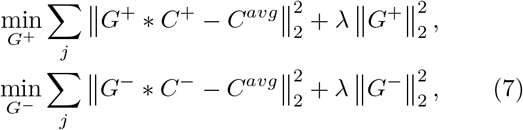

where *λ* is a Tikhonov regularization parameter controlling the tradeoff between noise amplification and bias. After training, the kernels are applied to correct the acquired time-series data:

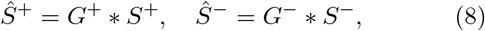

where *Ŝ*^+^ and *Ŝ*^−^ denote the corrected ghost-free RO+ and RO-data, respectively.

Notably, DPG uses a single kernel to correct one polarity, implicitly assuming that Φ_*err*_(**r, k**) is invariant along the *k*_*x*_ direction. This assumption can break down in demanding EPI trajectories that employ high gradient amplitudes, slew rates, and ramp sampling portion, and suffer from pronounced spatiotemporal phase variations.

### 2.4 Multi-kernel DPG (mk-DPG) for spatiotemporal varying phase errors along *k*_x_

To account for phase error variations along *k*_*x*_, we first developed multi-kernel DPG (mk-DPG) ^27^. Instead of training a single pair of kernels for the entire RO+ and RO-sets, in mk-DPG, the data along *k*_*x*_ are divided into multiple segments (Figure 1C). A corresponding pair of kernels is then trained for each segment:

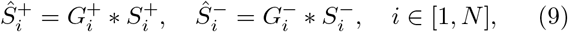

where *N* denotes the number of segments. When *N* = 1, mk-DPG converges to DPG. Figure 1C illustrates the mk-DPG workflow with *N* kernel pairs.

Increasing the number of kernels improves the ability to capture phase variations along *k*_*x*_. However, this comes at the cost of reduced training data per kernel (given a fixed amount of calibration data), which can degrade kernel estimation quality. Therefore, selecting an appropriate number of segments is critical for optimal mk-DPG performance.

### 2.5 Field Correcting GRAPPA (FCG): compact representation of *k*_*x*_-varying phase

Given the observation that Φ_*err*_ varies smoothly along *k*_*x*_, the corresponding kernel weights are expected to be highly correlated. To exploit such correlation, we developed Field-Correcting GRAPPA (FCG), which provides a compact and continuous representation of *k*_*x*_-dependent kernels. Unlike mk-DPG, where each kernel is independently trained and represented, FCG models the kernel as a function of *k*_*x*_ using a multi-layer perceptron (MLP)^28,29^. The network is trained jointly using all calibration data:

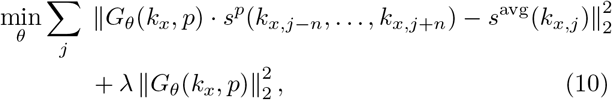

where *G*_*θ*_(*k*_*x*_, *p*) denotes the MLP parameterized by *ε*, and *p* indicates the readout polarity (RO+ or RO-). The input *s*^*p*^(*k*_*x,j*−*n*_,. .., *k*_*x,j*+*n*_) represents the source data centered at position *j* with *n* neighboring points on each side (yielding 2*n* + 1 samples), and *s*^avg^(*k*_*x,j*_) is the corresponding target data point. By traversing k-space, all source–target pairs are collected to train the network (Figure 1D). Compared with training multiple independent kernels, the shared representation in the MLP improves the conditions of the problem, and thus the SNR and robustness. Once the calibration data are acquired at the beginning of each scan, the training of *G*_*θ*_ is fully self-supervised.

## 3 METHODS

The studies in this work have three primary objectives:

- To develop the FCG technique with a flexible pipeline applicable to diverse EPI experiments.
- To validate the spatiotemporal phase error patterns during EPI sampling using gold-standard field probe measurements.
- To evaluate FCG across a range of experiments with varying gradient specifications, acceleration factors, and field strengths.

Detailed imaging protocols for all experiments are listed in Table 1. All in vivo studies were approved by the institutional review board, and written informed consent was obtained from all participants prior to scanning.

**TABLE 1.** Imaging protocols.

| Field Strength | Scanner | FOV (mm<sup>2</sup>) | In-plane res | Slice thickness | R | SMS factor | TR/TE (ms) | ESP (ms) | FA (°) |
| --- | --- | --- | --- | --- | --- | --- | --- | --- | --- |
| 3T | UHP | 220×220 | 1.1 mm | 2.0 mm | 4 | 1 | 2000/30 | 0.93 | 80 |
| 3T | UHP | 220×220 | 1.1 mm | 2.0 mm | 4 | 1 | 2000/30 | 0.73 | 80 |
| 3T | MAGNUS | 240×240 | 1.0 mm | 2.0 mm | 2 | 1 | 2000/30 | 0.58 | 80 |
| 3T | MAGNUS | 240×240 | 1.0 mm | 2.0 mm | 2 | 2 | 2000/30 | 0.58 | 80 |
| 7T | Impulse | 216×216 | 0.8 mm | 0.88 mm | 4 | 1 | 2100/25 | 0.60 | 70 |

### 3.1 Data correction pipeline for slice-by-slice and SMS EPI

In this work, DPG, mk-DPG, and FCG share the same data correction pipeline as shown in Figure 2A: (1) calibration data acquisition; (2) training data preparation; (3) kernel training; (4) imaging data correction. For slice-by-slice EPI, the step (1) (calibration data acquisition) and step (2) (training data preparation) are the same as specified in Section 2.3 and Figure 1. As a breif recap, in step (1), fully-sampled calibration data are acquired using the imaging EPI trajectories with dual polarities. In step (2), the calibration data are corrected using LGC first; the RO+ source data are formed by aggregating all RO+ readouts from the dual-polarity calibration data; the RO-source data are constructed analogously; the target data are obtained as the complex average of the RO+ and RO-source data. In step (3), kernel training is performed using the constructed source and target datasets for DPG, mk-DPG, or FCG. In step (4), the acquired time-series EPI data is corrected with LGC first, followed by correction of RO+ and RO-readouts using the trained kernels; this produces phase-error-corrected time-series EPI data that are good for following parallel-imaging reconstruction.

**FIGURE 2.**
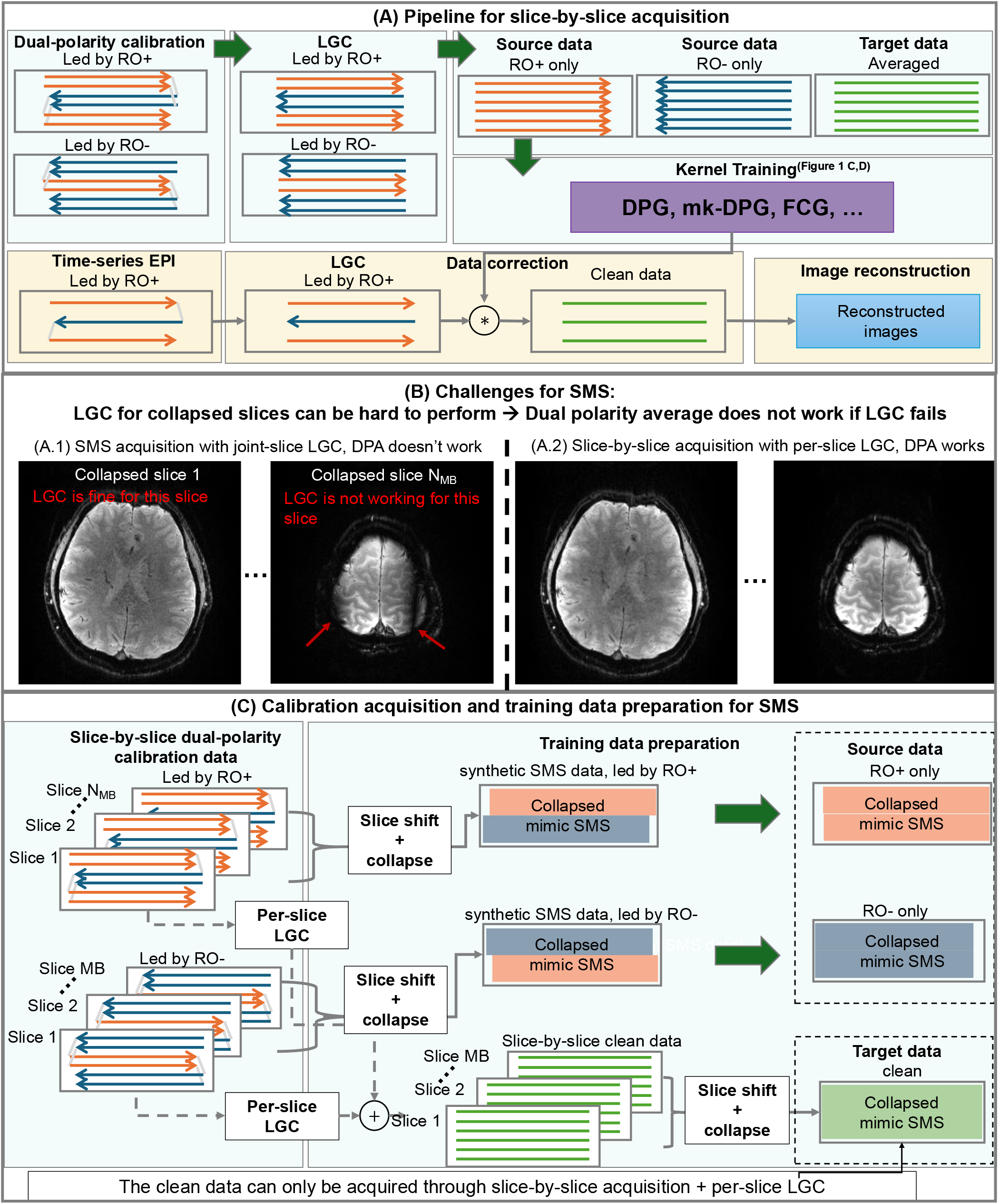
Fig 2. Data correction pipeline for slice-by-slice EPI and SMS-EPI. (A) Data correction pipeline for slice-by-slice EPI. The dual-polarity calibration data acquisition and source/target data preparation are the same as in Figure 1A and 1B. This kernel can be directly applied to time-series EPI. (B) challenges for SMS-EPI with strong phase errors. If the eddy-current-induced phase errors differ within the SMS slices, even DPA cannot provide clean results. There will be signal cancellation due to non-optimal LGC. (C) To resolve this issue, the calibration data for SMS is acquired in a slice-by-slice manner. The slice-by-slice calibration data are slice-shifted and collapsed as synthetic SMS source data. To generate the clean target data, the calibration data first receives optimal per-slice LGC for each and DPA to become clean slice-by-slice data. The clean slice-by-slice data is then slice-shifted and collapsed as synthetic clean target data. The kernels are then trained on the source and target data same as shown in Figure 1.

Application to SMS-EPI faces additional challenges. Slices at different spatial locations can have different linear components of phase errors. When calibration data are acquired directly with SMS manner, only a single set of LGC coefficients can be applied to simultaneously-excited slices, resulting in residual slice-dependent linear phase errors. These residual errors can lead to signal cancellation during dual-polarity averaging, as illustrated in Figure 2B. To address this issue, calibration data are instead acquired in a slice-by-slice manner. The source data are synthesized phase-error-corrupted SMS data by collapsing the uncorrected slice-by-slice calibration data. Note that if the blipped-CAIPI strategy is used in SMS acquisition to achieve improved undersampling efficiency ^24^, the synthesis of SMS data will include the spatial shift of the simultaneously-excited slices before collapsing them. To construct ghost-free target data, the slice-by-slice calibration data are corrected using slice-specific optimal LGC parameters, averaged across polarities to generate ghost-free slice-by-slice data, and then collapsed to form the corresponding ghost-free target SMS data using the same SMS synthesizing strategy as for the source data (Figure 2C).

### 3.2 FCG kernel training

With the acquired high-quality training data, FCG kernel training is straightforward. A large number of labeled training pairs are generated from the source and target datasets, as described in Section 3.1. For each training pair, the input consists of multi-channel k-space samples centered at an arbitrary location *j*, along with 2*n* neighboring samples spanning both positive and negative readout directions from the source data. The corresponding target is the multi-channel k-space data point at location *j* from the target dataset. For a typical 2D EPI dataset with matrix size 200 × 200, approximately 40,000 training pairs are available. The training process follows Equation 10. The MLP network consists of three fully connected layers, each with 256 neurons, and a hidden layer with 64 neurons. The Tikhonov regularization parameter *λ* in Equation 10 is by default set to 0.

Once trained, the network can be directly applied to correct the acquired EPI time-series data. For each multi-channel data point, the *k*_*x*_ location *j*, the corresponding neighborhood data points, and readout polarity *p* are fed into the network *G*_*θ*_ to produce a corrected multi-channel data point at that location. Applying this process across all k-space data points yields ghost-free EPI data.

### 3.3 Eddy-current phase characterization with field camera

Field cameras can provide gold-standard delineation of phase patterns during acquisition, but has limited accessibility. In this work, experiments with a standalone Skope system (Skope MR Technologies Inc., Zurich, Switzerland) were performed on a 3T whole-body scanner (UHP, GE Healthcare, Milwaukee, USA) to evaluate the phase errors and validate the following assumptions: (1) dual-polarity averaging (DPA) can effectively suppress odd–even phase errors, which supports DPA, DPG, and FCG; and (2) phase errors vary along *k*_*x*_ but remain relatively stable along *k*_*y*_, a key assumption in the current FCG framework.

Two EPI trajectories were designed for the study. The first one is a standard trajectory with FOV = 220×220 mm^2^, in-plane resolution = 1.1×1.1 mm^2^, maximum gradient amplitude (gmax) = 40 mT/m, maximum slew rate (smax) = 120 T/m/s, ramp sampling factor = 64%, echo spacing (ESP) = 0.93 ms, and undersampling factor *R* = 4. The gmax and smax of a standard EPI trajectories are usually smaller than the hardware maximum (which is 80 mT/m and 200 T/m/s for gradient amplitude and slew rate, respectively, for a UHP system) due to the limit of peripheral nerve stimulation (PNS). The second trajectory utilizes more aggressive gradient performance for faster sampling at the same res-olution: gmax = 50 mT/m, smax = 200 T/m/s, ramp sampling factor = 68%, ESP = 0.73 ms, and *R* = 4. Phase perturbations for all interleaves were measured using Skope.

The measured phase data were analyzed in three aspects: (1) the magnitude of odd–even phase errors; (2) variation of phase errors along *k*_*x*_ and *k*_*y*_; and (3) residual phase errors after dual-polarity averaging.

Phantom and in vivo data were acquired on the same system using a commercial 32-channel head coil (NOVA Medical, Wilmington, MA) with a gradient echo (GRE) sequence. The dual-polarity calibration data (for phase-error correction and parallel imaging reconstruction) were acquired first, followed by time-series EPI data (*R* = 4) acquired with alternating polarity to enable DPA reconstruction as a reference. Twenty repetitions were collected for temporal SNR (tSNR) analysis. The following reconstruction methods were compared: (1) expanded encoding model using Skope-measured phase maps ^30,31^; (2) DPA of two consecutive EPI shots with alternating polarity followed by GRAPPA; and (3) DPG, mk-DPG, and FCG applied to single-average data, followed by GRAPPA. Details of reconstruction using Skope-measured phases are provided in Supporting Information Section B.

### 3.4 Fast EPI sampling on high-performance 3T head-only system

To evaluate FCG under challenging conditions with rapid gradient switching, phantom and in vivo experiments were conducted on a high-performance 3T head-only system (SIGNA™ MAGNUS, GE Healthcare, Milwaukee, USA, with maximum gradient amplitude of 300 mT/m and slew rate of 750 T/m/s using standard GE Health-Care 3.0 T system power electronics of 2 MVA peak driver power per axis). A GRE-EPI sequence was used with the following parameters: FOV = 240 × 240 mm^2^, in-plane resolution = 1.0 mm^2^, gmax = 50 mT/m, smax = 600 T/m/s, ESP = 0.58 ms, ramp sampling factor = 29%, and *R* = 2. The readout gmax used for the EPI trajectories is limited by the sampling dwell time of the ADC (2*µs*). Given this limit, a gmax larger than 50 mT/m can lead to undersampling in the readout direction.

Two acquisition schemes were conducted: (1) slice-by-slice EPI and (2) SMS-EPI with SMS factor = 2 and 40-mm separation. The data acquisition process is the same as in Section 3.3: dual-polarity calibration data are acquired first, followed by time-series EPI data with alternating polarity over 20 repetitions. The imaging data were corrected using DPA, DPG, mk-DPG, and FCG, followed by in-plane GRAPPA and/or slice-GRAPPA reconstruction.

### 3.5 High-resolution EPI application on head-only system at high field

To evaluate FCG at higher field strength, in vivo experiments were performed on a 7T head-only system (Impulse, Siemens Healthineers, Germany, with maximum gradient amplitude of 200 mT/m and slew rate of 900 T/m/s) with a 8Tx/64Rx channel head coil (MR CoilTech, Glasgow, UK). A in-house GRE-EPI sequence was used with the following parameters: FOV = 216×216 mm^2^, in-plane resolution = 0.8 mm^2^, *R* = 4, gmax = 55 mT/m, smax = 750 T/m/s, ramp sampling factor = 25%, and ESP = 0.60 ms. The acquisition protocol is the same as described in Section 3.4.

### 3.6 Data analysis

All image reconstruction and data processing were performed in Python using an NVIDIA RTX A5000 GPU. In all experiments, DPA was set as the reference. The image quality using DPG, mk-DPG, and FCG were evaluated by quantifying their relative root mean squared error (rRMSE) against DPA. To characterize mk-DPG performance, rRMSE was evaluated across a range of *k*_*x*_ segmentation numbers (*N* = 2 to 20 with increment of to 1) determine the optimal number of segmentation. The tSNR^32^ was also evaluated, which is particularly relevant for applications in fMRI. The tSNR maps were computed following the standard definition ^33^ as the ratio of the temporal mean to the temporal standard deviation. For DPG, mk-DPG, and FCG, only data led by the same polarity (e.g., led by RO+) were used for tSNR calculation. For DPA, since data are averaged across two consecutive TRs / shots, the resulting tSNR was scaled by 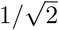 for fair comparison.

For all the phase error correction approaches in this work, Tikhonov regularization controls the trade-off between noise amplification and reconstruction bias. Although the *λ* was set to 0 by default, it is important to understand their reaction to regularization for different approaches. To evaluate this effect, rRMSE and tSNR were computed across a range of *λ* values for DPG, mk-DPG (Equation 7), and FCG (Equation 10). For DPG, *λ* values ranging from 0 to 10^−2^ (step size 10^−3^) were tested. For mk-DPG, the regularization parameter was scaled as *λ*_mk_ = *λ*_DPG_*/N* given *N* segments because the data consistency term in Equation 7 is reduced by N. For FCG, *λ*_FCG_ was scaled relative to *λ*_DPG_ to ensure comparable rRMSE ranges. Convergence of FCG training was also evaluated as a function of training epochs.

## 4 RESULTS

### 4.1 Characterization of phase errors with Skope

Figure 3 shows the spatiotemporal phase perturbations measured by Skope. The readout gradient *G*_*x*_ for the regular and fast EPI trajectories, along with their corresponding 0-order phase coefficients, are displayed in Figure 3A. During EPI encoding, the 0-order phase exhibits two components: a smooth global drifting along *k*_*y*_, and a rapidly varying component associated with *G*_*x*_ switching. The average phase difference between adjacent RO+ and RO-is 0.62 radians for the regular trajectory and 0.89 radians for the fast trajectory, indicating stronger eddy-current effects caused by fast gradient switching.

**FIGURE 3.**
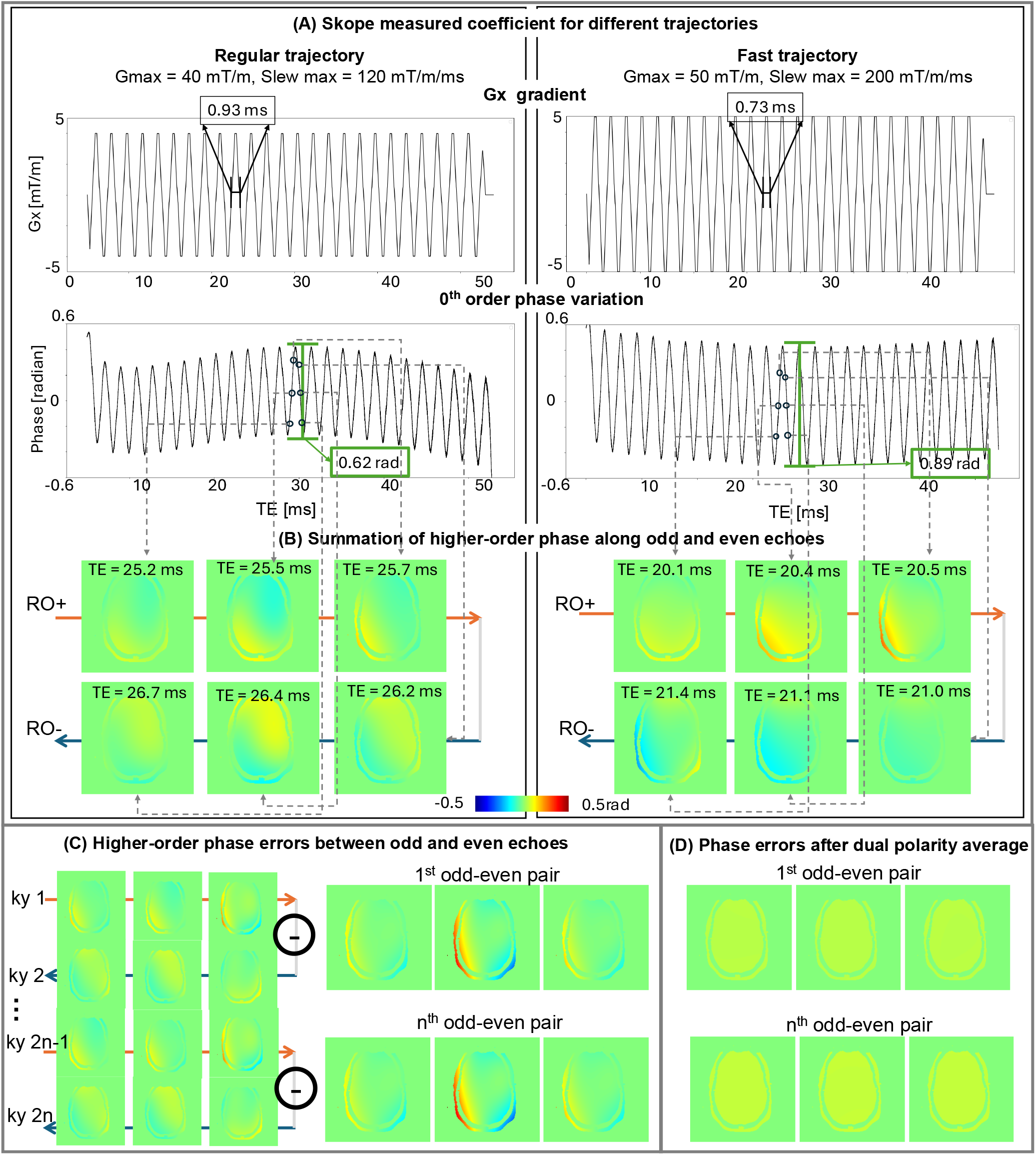
Skope measurement of the phase variation during EPI encoding. (A) Two EPI trajectories are measured. One is a regular trajectory with medium gradient and slew rate. The other is a fast trajectory with slew rate close to the upper limit of the system. The faster trajectory showed larger phase variation between RO+ and RO-. (B) The odd-even phase errors between RO+ and RO-vary along kx. (C) The odd-even phase errors stay consistent along ky (although the absolute phase pattern along ky may have smooth variation, as shown in A). (D) When averaging the odd-even phase errors between trajectories with opposing polarity, the phase errors are mostly cancelled out. This supports the success of DPA.

The total phase perturbation maps (excluding the first-order term) are shown in Figure 3B at six time points near the *k*_*y*_ = 0. The fast trajectory exhibits substantially larger phase perturbations. The phase errors between adjacent RO+ and RO-lines (Figure 3C) vary along *k*_*x*_ but remain stable along *k*_*y*_, which consolidates a key assumption for mk-DPG and FCG. Note that the *B*_0_ induced phase was not included in the analysis.

After averaging the phase patterns measured with trajectories led by RO+ and RO-, the residual phase differences between adjacent RO+ and RO-lines are reduced to below 0.1 radians, confirming that DPA effectively suppresses eddy-current-induced phase errors (Figure 3D).

In the reconstruction, the Skope-based reconstruction is overall consistent with the DPA references, with minor differences primarily at the phantom edges (Figure 4), which are likely distortions from the global phase drift in the Skope measurement, as illustrated in the Supporting Information Figure S1. For the comparison of LGC, DPG, mk-DPG, and FCG, LGC leaves strong ghosting and shading artifacts; DPG substantially reduces these artifacts, although residual ghosting remains; mk-DPG, with an optimal configuration of five kernel pairs, further improves the results; FCG achieves the lowest error, with an rRMSE of 1.1%. The dependence of mk-DPG’s performance on the number of kernel pairs is provided in Supporting Information Figure S2.

**FIGURE 4.**
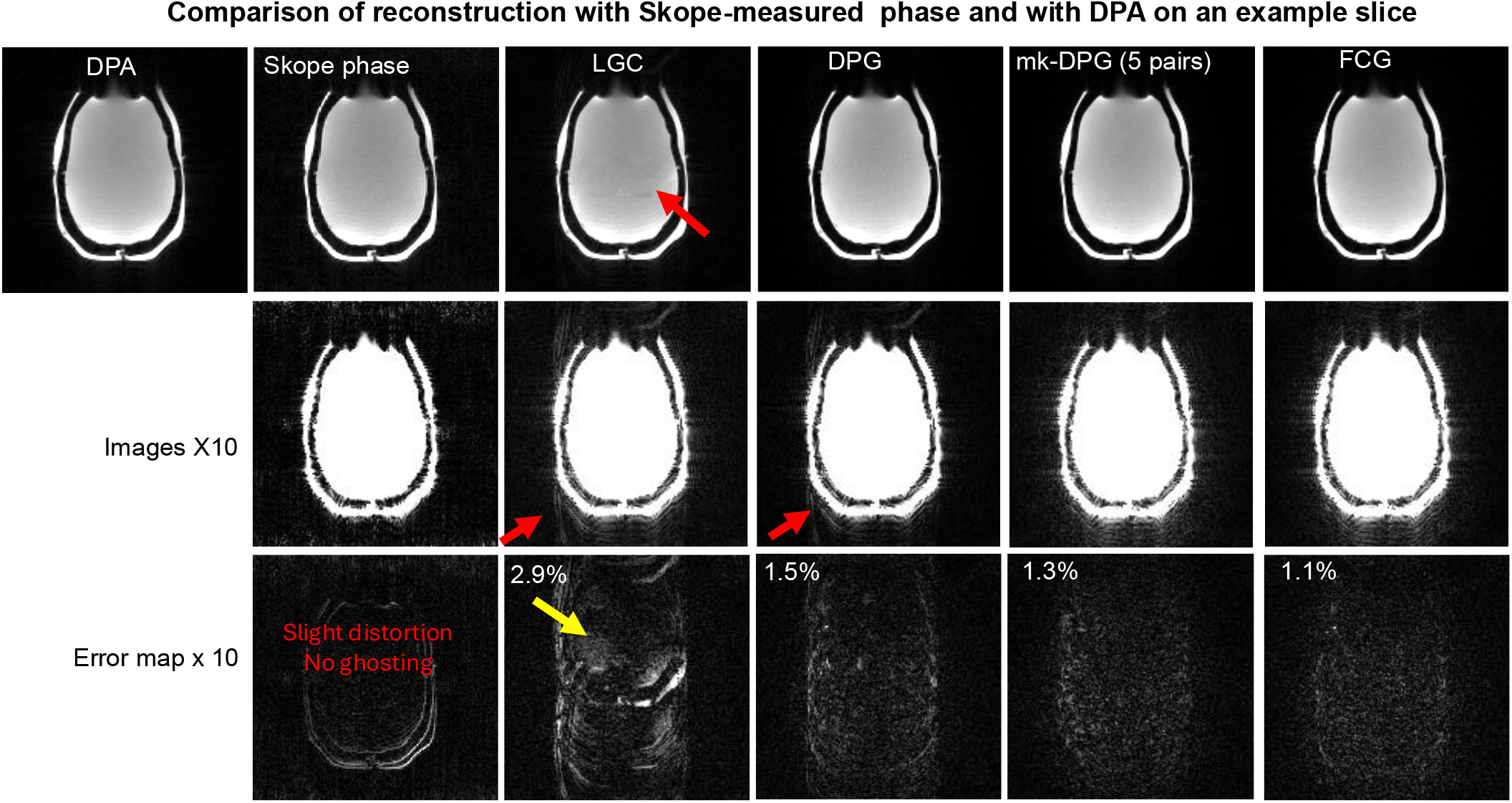
Comparison of reconstruction with Skope measured phases vs DPA. Both methods were able to remove eddy-current-induced artifacts in EPI. The differences between DPA and Skope-based reconstruction are mainly in the edges of the structure. Such differences in the distortion come from the smooth global phase variation in Skope measurement, which works similarly to the B0 phase accumulation. FCG shows the lowest error compared to DPA reference. The red arrows in the figure indicate the aliasing artifacts, and the yellow arrows point to the shading artifacts.

### 4.2 Fast EPI sampling on high-performance 3T head-only system

Figure 5 demonstrates the performance under fast EPI sampling with high slew rate and short echo spacing. Observations are similar. With LGC only, severe ghosting is observed. DPG improves image quality but leaves noticeable residual artifacts. mk-DPG further reduces these artifacts but suffers from increased noise. The optimal number of kernel sets in this experiment is seven, higher than five observed in Section 4.1, suggesting stronger phase variation along *k*_x_ . In contrast, FCG produces ghost-free images while maintaining low noise levels.

**FIGURE 5.**
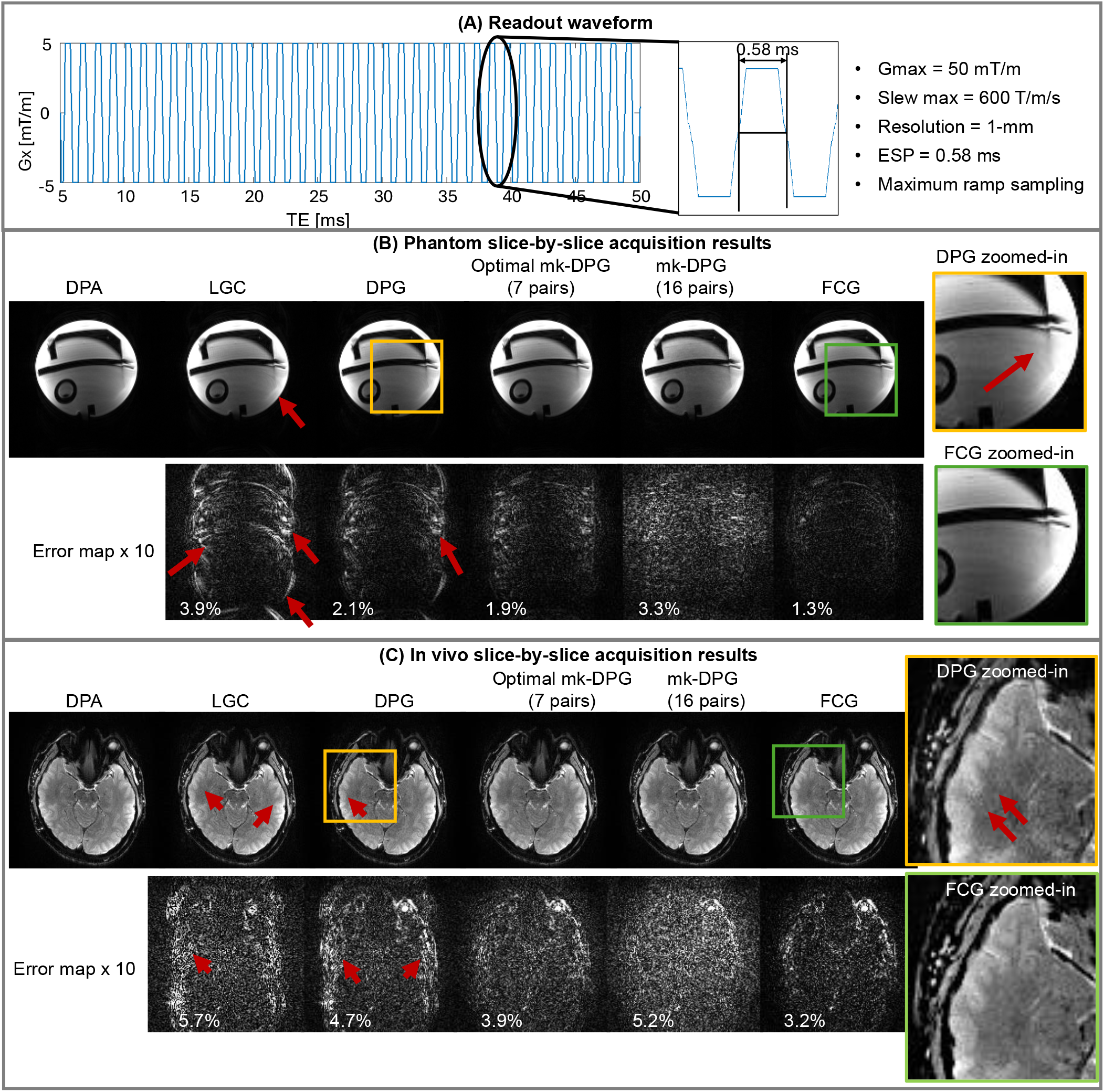
The phantom and in vivo results from 3T head-only MAGNUS system. (A) The EPI trajectory used on MAGNUS. Ultra high slew rate and maximum ramp sampling are used to create a challenging case for eddy current correction. (B) The phantom results (C) The in vivo results. In (B) and (C), the red arrows indicate the ghosting artifacts. FCG produced the lowest error against DPA for both phantom and in vivo studies.

In the SMS experiment, simultaneously acquired slices separated by 40 mm exhibit substantially different linear phase errors (Figure 6A). Under these conditions, direct application of DPA of SMS data fails to produce clean reconstructions. As shown in the “DPA-SMS” column of Figure 6B–C, one or more slices exhibit shading and signal cancellation. In contrast, DPG, mk-DPG, and FCG that are trained on synthetic data from slice-by-slice calibration scans substantially reduce these artifacts. Among them, FCG provides the best performance with minimal ghosting and low noise.

**FIGURE 6.**
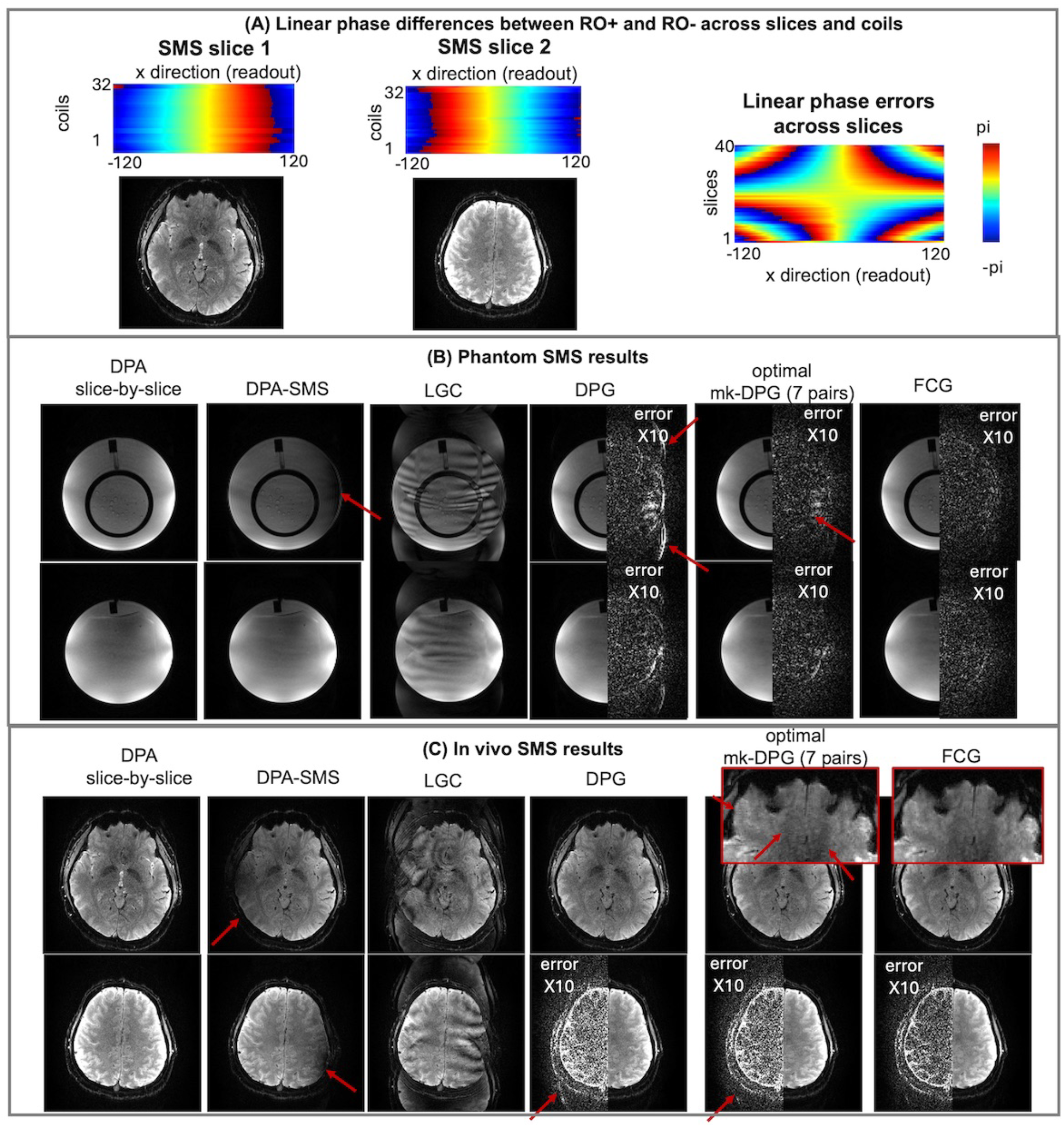
The SMS results from 3T head-only MAGNUS system. (A) when acquiring the data in slice-by-slice manner and estimate the LGC per slices, the linear component of phase difference vary a lot across slices, increasing the difficulties for SMS correction. In both phantom (B) and in vivo (C) studies, directly DPA of SMS cannot provide a clean output due to the LGC for the collapsed slices not being optimal. For DPG, mk-DPG, and FCG, whose kernels are trained on the slice-by-slice calibration data, the results are highly improved, with FCG producing the highest image quality and lowest noise level. The red arrows indicate signal cancellations, ghosting artifacts, or noise.

The residual differences between FCG and slice-by-slice DPA are primarily localized at tissue boundaries for the in vivo experiments (Figure 6C), which are likely caused by minimal residual bulk motion between calibration and SMS acquisitions, or within the dual-polarity calibration data. Nevertheless, because the phase errors are spatially smooth, kernels learned from calibration data are generalized well to SMS data despite such motion.

### 4.3 High-resolution EPI on head-only system at high field

Figure 7 demonstrates the performance of FCG on the 7T head-only system. Under challenging conditions with high slew rate, high spatial resolution, and high in-plane acceleration (R=4), FCG achieves nearly ghost-free reconstruction comparable to DPA. At the same time, FCG yields the highest tSNR. Note that tSNR of DPA is scaled by a factor of 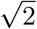 for fair comparison. The voxel-wise tSNR difference histograms (Figure 8) show that FCG outperforms both DPG and optimally-tuned mk-DPG in the majority of voxels.

**FIGURE 7.**
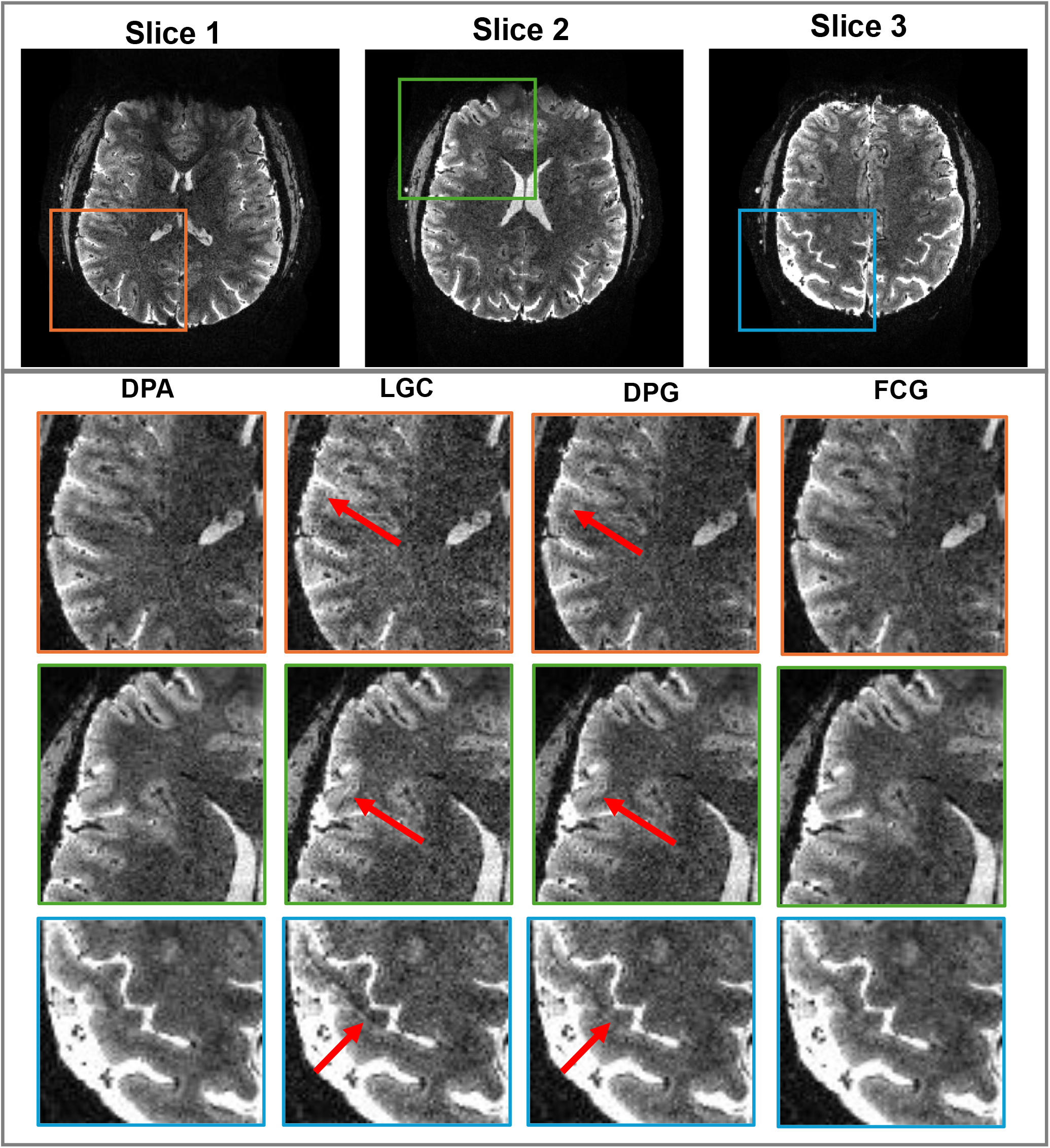
The 0.8-mm isotropic EPI study on 7T Impulse system. For three representative slices, the single-average image shows strong residual artifacts, which cannot be resolved by DPG. In this case, FCG consistently produces clean output against DPA.

**FIGURE 8.**
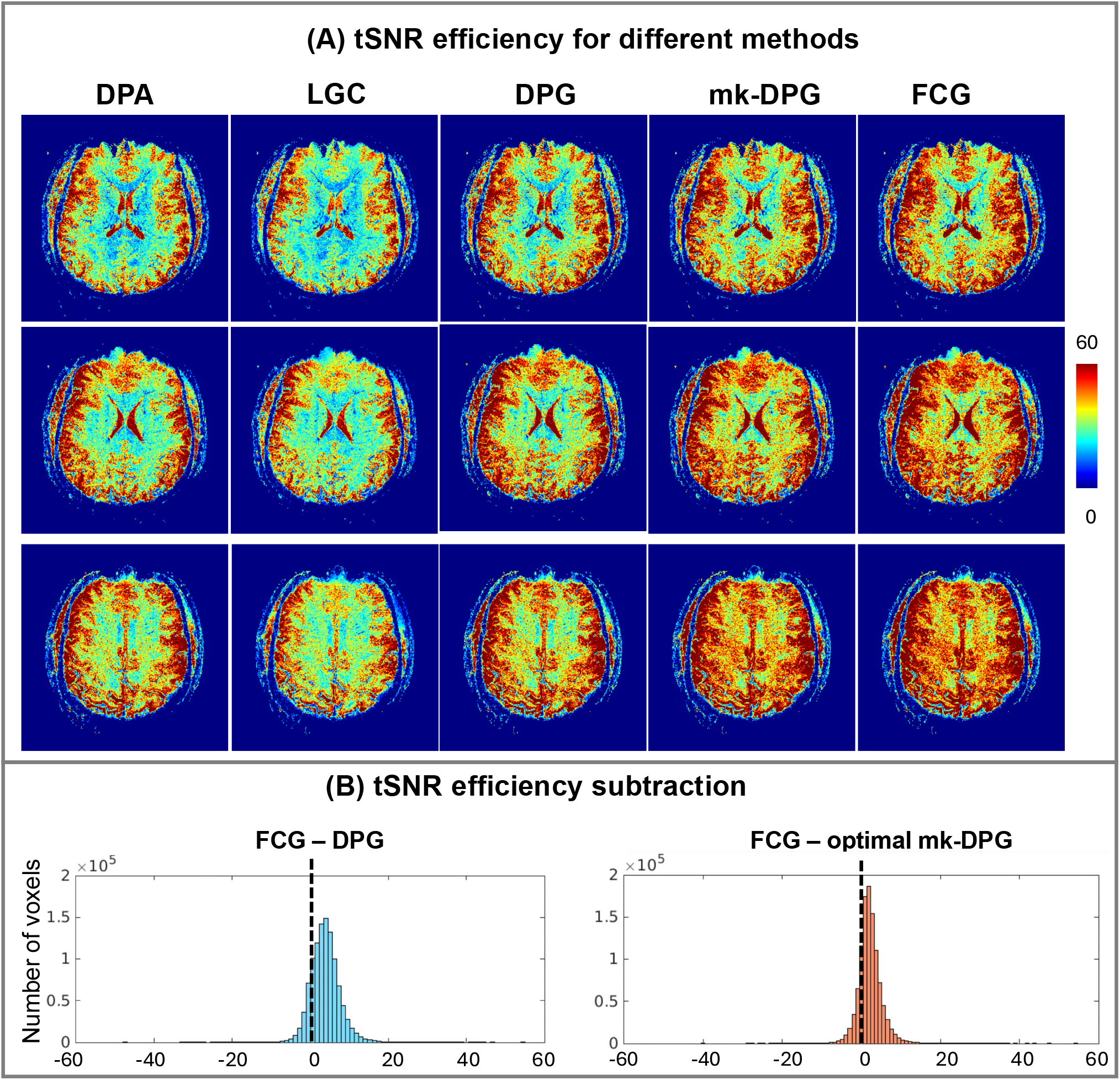
The tSNR efficiency for DPA, single-average, DPG, mk-DPG, and FCG on the 7T Impulse data. FCG shows the highest tSNR efficiency. (B) The histogram of the tSNR efficiency subtraction for FCG vs DPG and FCG vs mk-DPG. For most of the voxels, FCG bears the higher tSNR efficiency against DPG or mk-DPG.

### 4.4 Robustness of FCG

The MLP used in FCG is compact and converges rapidly. As shown in Figure 9A, the rRMSE stabilizes within 10 epochs. Each epoch requires about 0.3 seconds on the GPU, resulting in a total training time of 3–6 seconds per slice. For comparison, DPG kernel training requires approximately 1 second per slice. FCG also demonstrates robustness across a wide range of Tikhonov regularization parameters. L-curve analysis, showing the trade-off between noise (quantified as 1*/*tSNR) and reconstruction bias (rRMSE relative to DPA), indicates that FCG consistently operates the best.

**FIGURE 9.**
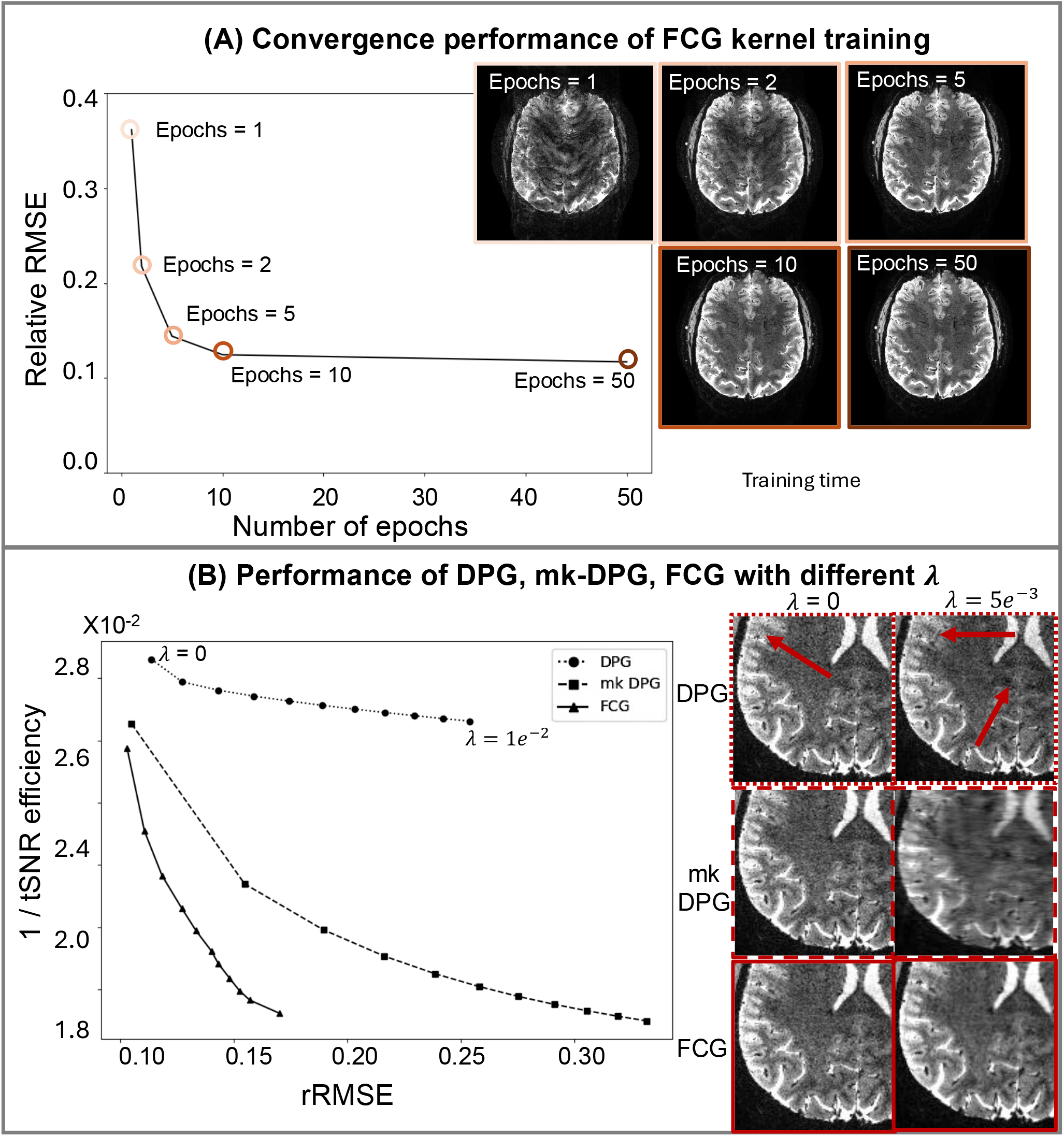
The robustness of FCG. (A) The convergence performance of MLP kernels with different numbers of epochs. (B) The L-curves show the trade-off between noise (1/tSNR) and reconstruction error (rRMSE relative to DPA). FCG consistently provides the lowst error and lowest noise level at different regularization level compared to DPG and ml-DPG (with optimal number of kernel pairs, which is 5 in this example).

## 5 DISCUSSIONS

Eddy-current-related artifacts has been a long-standing problem in MRI, with different appearances for different sampling strategies. In the regime of EPI, the fast-changing readout gradient waveforms tend to produce spatiotemporal varying non-linear phase perturbations. The inconsistency of the phase perturbations between RO+ and RO-readouts are a major source of artifacts for EPI.

A category of existing techniques represent the phase errors, explicitly or implicitly, as a non-linear phase map between the RO+ and RO-, estimate such a map, and account for it during reconstruction. Example techniques include Chen and Wyrwicz ^34^, PAGE^35^, real-time PAGE^36^, GESTE^37^, and DPG^21^. An implicit assumption for these methods is that phase error map between RO+ and RO-is constant along *k*_*x*_. Another category is to directly average RO+ and RO-read-outs from dual polarity acquisitions, including double-sampled EPI^17^, PLACE^18^, and DPA^20,19^. These methods correct the acquired data point-by-point along *k*_*x*_, and therefore are capable of handling phase error variations along *k*_*x*_, However, they suffer from a double scan time.

The technique developed in this work, FCG, is able to account for the *k*_*x*_-varying phase errors without increasing the scan time. The key steps for FCG include: (1) obtaining a set of calibration data producing eddycurrent-corrupted data and clean data; (2) training a set of kernels representing the spatiotemporal-varying phase errors along *k*_*x*_; (3) cleaning the corrupted time-series data by applying the kernels to it. There are multi-fold assumptions that are crucial to the success of FCG: (1) averaged dual-polarity data can serve as ghost-free target data; (2) the odd-even phase errors vary along *k*_*x*_, but stay stable along *k*_*y*_. These assumptions were validated by the Skope measurement performed on UHP system. Figure 3D demonstrates that averaging dual-polarity data can largely suppress the odd-even phase errors; Figure 3C illustrates that the odd-even phase errors vary along *k*_*x*_, but stay stable along *k*_*y*_. Additionally, Skope measurement shows that EPI trajectories with higher slew rate produce elevated spatiotemporal phase variations (Figure 3B), which poses the need for FCG over DPG to reduce the ghosting artifacts in demanding EPI trajectories.

An interesting observation from Skope measurements is that, while the phase differences between adjacent RO+ and RO-echoes remain stable, the overall phases slowly and smoothly drift along echo train. This phase accumulation does not add on to the odd-even artifacts, but poses minor distortion. DPA cannot reduce this effect because the the signs of the phase drift are the same for dual polarity data, as shown in Figure S1. However, when reconstructing the images with Skope-measured phases, this distortion effect can be corrected. This explains the differences at the edge of the phantom images comparing DPA correction and reconstruction with Skope-measured phases.

The experiments on the 3T MAGNUS system demonstrated the potential of FCG to effectively remove the ghosting artifacts in demanding EPI trajectories. The EPI trajectory is an in-house design with gmax = 50 mT/m and smax = 600 T/m/s, with maximum ramp sampling portion and minimal gap between echoes. The resulting eddy-current-related phase errors produced significant amount of artifacts in the images even after convetional LGC. DPG correction improved the image quality, but still left substantial amount of errors. Both mk-DPG and FCG were able to remove the ghosting and shading. However, mk-DPG produced elevated amount of noise due to the reduced amount of training data for each pair of kernels. The pair of kernel sets may need to be tuned case by case depending on the image resolution and level of phase error variation. Moving to higher-resolution applications with faster slew or larger ramp sampling factor, a larger number of kernel sets will likely be needed to capture the larger spatiotemporal phase variations. On the other hand, FCG, as a compact representation of the spatiotemporal varying phase errors, consistently provided high-quality high-SNR images. In this work, the hyperparameters of the MLP network, including the number of layers, the size of each layer, and *λ* for Tikhonov level were kept consistent for all experiments, indicating its robustness to different EPI setups and system performance. In this study, the read-out gmax was limited to 50 mT/m because of a software limit of the sampling dwell time of ADC, which is 2 *µs*. If 1-*µs* sampling can be unlocked, the readout gmax can be increased to 100 mT/m, allowing further accelerated sampling.

The kernel learns a multi-channel representation of the nonlinear phase errors along *k*_*x*_. Given that the multi-channel coil sensitivity and the nonlinear phases are spatially smooth, the kernel is to some extent robust to small motions as long as support of the object has not significantly changed. For example, the in vivo case in Figure 6 bears a small displacement between the slice-by-slice calibration data and the actual time-series SMS imaging data, which is indicated by the elevated errors at the edge of the structures. In this scenario, the kernel trained on the calibration data can still produce high-quality correction on the imaging data. The kernel can be applied to imaging data with different contrast as long as the EPI trajectory is the same, e.g., to images with different b values in diffusion imaging.

The Skope measurement was not available for 3T MAGNUS system and 7T Impulse system. The two head-only systems have different hardware design ^38^ compared to whole-body UHP system, which can potentially produce different eddy-current patterns. The variation of phase errors can be, possibly, not only along readout, but also along echo train or through shots. In such cases, FCG can also be modified to capture multi-fold variations. In the future, more characterizations of the phase perturbations on these systems can improve the understanding of phase perturbations as well as the development of FCG.

FCG can potentially benefit many EPI applications. Huber et al. showed that DPA has been a robust solution to eddy-current-induced fuzzy-ripple artifacts in fMRI^20^, which is a significant limitation for pursuing high-resolution protocol. Stirnberg et al. ^19^ illustrated that DPA can significantly improve the quality of QSM and R2* mapping with EPI. Investigations in Jalnefjord et al ^39^. exhibited that DPA can improve the diffusion MRI with EPI. In these applications, and potentially other applications, FCG can be a promising technique to remove the eddy-current-induced artifacts without the need for acquiring all data in dual polarity, which maintains the image quality while keeping the total scan time. The pipeline to apply FCG is straightforward and allows easy clinical and scientific translational practice.

This work has several limitations. First, the dual polarity calibration data are acquired with slices as the inner loop, and polarities as the outer loop. For in vivo study, respiration, cardiac pulsation, and bulk motion can potentially introduce shot-to-shot signal variation and thus can reduce the quality of calibration data when the dual-polarity shots are acquired far apart. In the future, FLEET structure ^40^, which acquires all the shots of one plane in a continuous manner with modified flip angle, will be implemented for calibration acquisition to improve the robustness of the calibration data. A second limitation is that the kernel only models the phase errors variation along *k*_*x*_. For future studies on more demanding EPI trajectories on high-performance head-only systems, characteristic of the eddy current patterns and modifications on FCG network to include variation along *k*_*y*_ can be helpful.

## 6 CONCLUSIONS

In this work, we developed FCG, which effectively corrects the spatiotemporal varying eddy current artifacts and produces high-quality EPI images. The performance of FCG is demonstrated to be superior across multiple experiments with demanding EPI trajectories on high-performance systems at different field strengths. The acquisition and reconstruction pipeline is straightforward and flexible for different EPI applications including SMS acquisition.

## ACKNOWLEDGMENTS

Authors HM and OS are supported by Deutsche Forschungs Gemeinschaft (DFG), Project Nr. 447709127

## Author contributions

Authors NW and DA contributed equally to this work.

## SUPPORTING INFORMATION

The following supporting information is available as part of the online article:

**Figure S1**. (A) Skope measured temporal coefficient of the phase perturbations. (B) The difference between the first-order temporal coefficient and the nominal trajectories.

**Figure S2**. Performance of mk-DPG with different pairs of kernels

