## Supplementary material for "Field-Correcting GRAPPA (FCG): a technique to correct spatiotemporal-varying phase errors in Echo Planar Imaging": figure S1 and S2

A. **Cancellation of B0’s effect along readout direction in dual-polarity averaging**

When including the effect of B0, Equation 5 is expended as:

$$S^{+}\left( \mathbf{k} \right)= \int_{\mathbf{r}} M_{xy}\left( \mathbf{r} \right)e^{i\mathbf{kr}+i\Phi_{ec}\left( \mathbf{r,k} \right)+iB0\left( \mathbf{r} \right)T(k_{x})+iB0\left( \mathbf{r} \right)T(k_{y}+k_{z})}d\mathbf{r},$$

$$S^{-}\left( \mathbf{k} \right)= \int_{\mathbf{r}} M_{xy}(\mathbf{r})e^{i\mathbf{kr}-i\Phi_{ec}\left( \mathbf{r,k} \right)-iB0\left( \mathbf{r} \right)T(k_{x})+iB0\left( \mathbf{r} \right)T(k_{y}+k_{z})}d\mathbf{r}. (S.1)$$

where $T(k_{x})$ is the function representing time t as a function of k-space location $\mathbf{k}$. For a k-space location ${(k}_{x},k_{y},k_{z})$ acquired with alternating polarity, the B0 phase accumulations along phase encoding direction are the same, but along readouts antisymmetric. Averaging $S^{+}\left( \mathbf{k} \right)$ and $S^{-}\left( \mathbf{k} \right)$ yields

$$S^{\mathrm{avg}}\left( \mathbf{k} \right)= \int_{\mathbf{r}} M_{xy}\left( \mathbf{r} \right){\cos\left( \Phi_{ec}\left( \mathbf{r,k} \right) \right)\cos\left( B0\left( \mathbf{r} \right)T(k_{x}) \right)e}^{i\mathbf{kr}+iB0\left( \mathbf{r} \right)T(k_{y}+k_{z})}d\mathbf{r}. (S.2)$$

Since the echo spacing in EPI is usually less than 2 ms, the phase accumulation along readout $B0\left( \mathbf{r} \right)T(k_{x})$ is small and smooth, resulting in $\cos\left( B0\left( \mathbf{r} \right)T(k_{x}) \right)\approx1$. This indicates that DPA can also suppress the image artifacts from the phase errors accumulated along readouts due to B0.

B. **Expanded encoding model for the reconstruction with Skope-measured phase perturbations**

As shown in Equation 2, the Skope measures the spatiotemporal phases during the performance of gradient waveforms and decomposes them into third-order spherical harmonic basis functions and the corresponding time-varying expansion coefficients with $\Phi\left( \mathbf{r},t \right)=\sum_{l=0}^{15} c_{l}(t)h_{l}(\mathbf{r})$, where $h_{l}$ is the spherical harmonic coefficients up to third order, and $c_{l}(t)$ is the temporal coefficient sampled at 1MHz frequency. The first-order temporal coefficient of x and y can be considered as the measured sampling trajectory $\tilde{k_{x}},$ $\tilde{k_{y}}$. They are likely to differ from nominal trajectory $k_{x}$ and $k_{y}$. Figure S1 shows example 0^th^ – 3^rd^ order temporal coefficients measured by Skope. Subtracting the measured trajectory $\tilde{k_{x}}$ and $\tilde{k_{y}}$ from the nominal trajectory $k_{x}$ and $k_{y}$, the residual differences are illustrated in Figure S1 (B).

To take the spatiotemporal phase errors into reconstruction without blurring it in the spatial or temporal dimension, an Expanded Encoding Model with conjugate gradient approximation has been used. It directly defines the imaging problem as

$$\sigma=E\rho, (S.3)$$

Where $\rho\in\mathbb{C}^{{N_{T}N}_{s}\times1}$, is the acquired data vector in size with $N_{T}$ as the total number of data (time) points acquired, $N_{s}$ is the total number of channels, $E\mathbb{\in C}^{{{N_{r}\times N}_{T}N}_{s}}$ is the encoding matrix, with $N_{r}$ as the total spatial points of the images. Each element in $E$ can be built as $E_{\left( t,v \right),\mathbf{r}}=S_{v}(\mathbf{r})e^{i\Phi\left( \mathbf{r},t \right)}$, without taking the B0-induced phase into consideration.

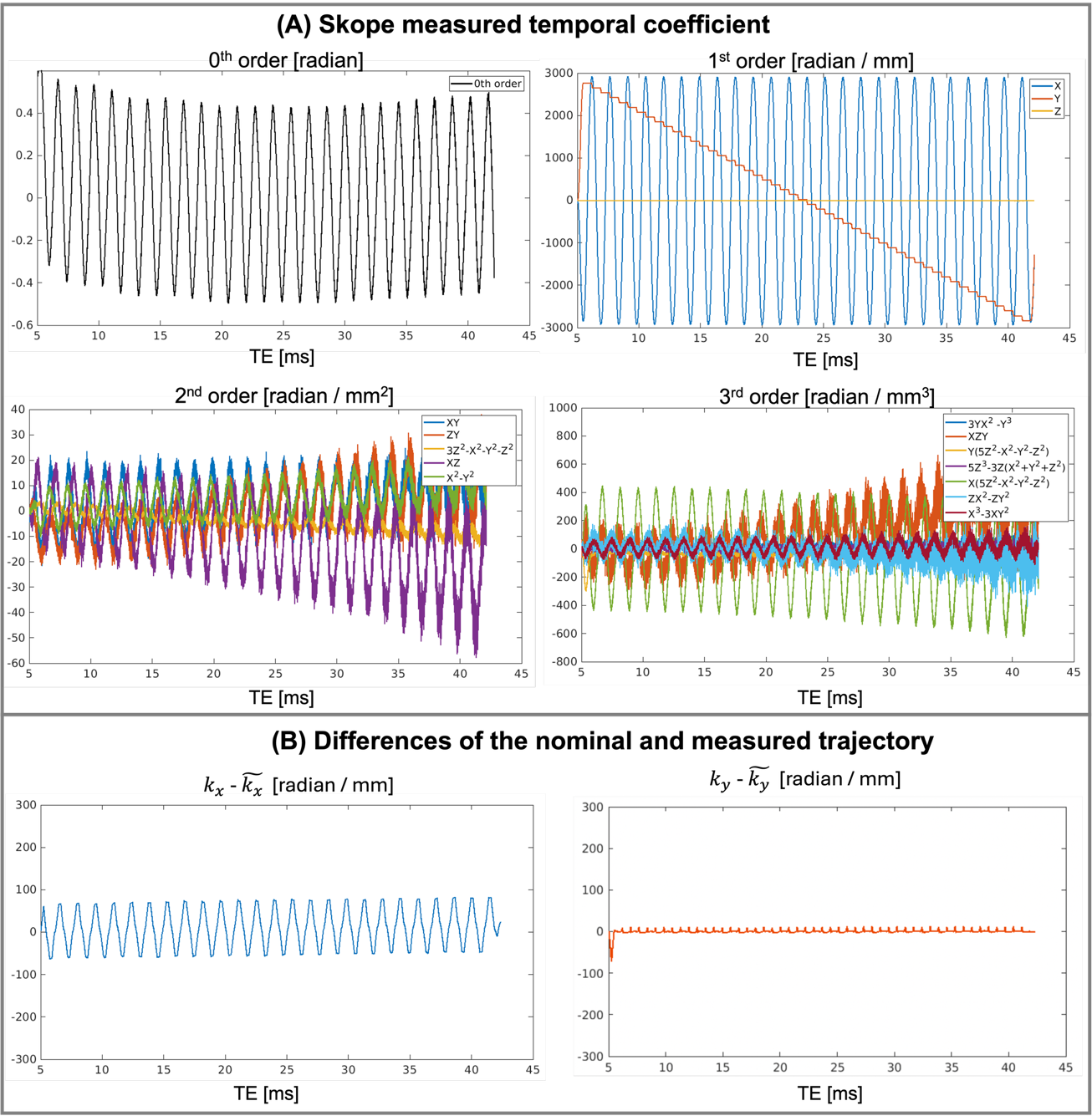

Figure S1: (A) Skope measured temporal coefficient of the phase perturbations. (B) The difference between the first-order temporal coefficient and the nominal trajectories.

The matrix $E$ is very expensive, and the direct reconstruction can be impractically memory-consuming. In ref 25, an algorithm with conjugate gradient approximation was developed, bringing the memory and time of the Expanded Encoding Model to a practical level. The matrix size of the Skope reconstruction is: $N_{r}=40000 (200 \times200)$, $N_{T}=20000$, $N_{s}=16$ (after SVD coil compression).

C. **Evaluation of mk-DPG**

To determine the best number of segments for mk-DPG, the rRMSE of mk-DPG was evaluated from N = 2 to 20 at an increment of 1. Figure S2 displays the rRMSE curve against N for the EPI experiments at 3T UHP, 3T Magnus, and 7T Impulse. The optimal number of segments varies for different experiments. The possible reasons are: (1) the amount of available training data (depending on the coverage and resolution of the data); (2) different eddy current patterns from different systems; (3) different levels of phase variation along k_x_ due to different resolution, slew rate, and ramp sampling portions. Therefore, although mk-DPG outperforms conventional DPG, it requires fine-tuning for each case to find the optimal number of kernels.

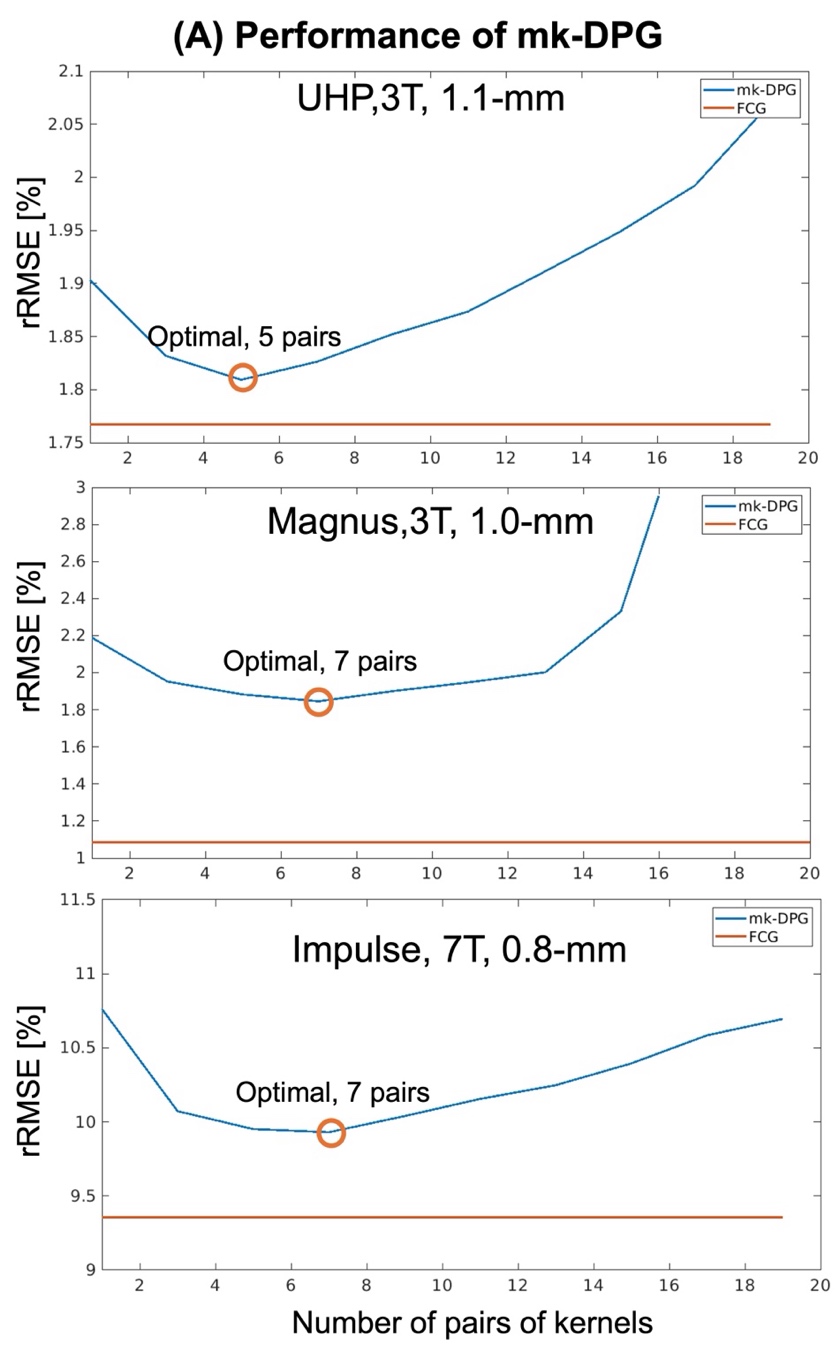

Figure S2: Performance of mk-DPG with different pairs of kernels
